# Simultaneous recording of ultrasonic vocalizations and sniffing from socially interacting individual rats using a miniature microphone

**DOI:** 10.1101/2023.09.02.556068

**Authors:** Shanah Rachel John, Rishika Tiwari, Yizhaq Goussha, Rotem Amar, Alex Bizer, Shai Netser, Shlomo Wagner

## Abstract

Vocalizations are pivotal in mammalian communication, especially in humans. Rodents accordingly rely on ultrasonic vocalizations (USVs) that reflect their internal state as a primary channel during social interactions. However, attributing vocalizations to specific individuals remains challenging, impeding internal state assessment. Rats emit 50 kHz USVs to indicate positive states and intensify sniffing during alertness and social interactions. Here, we present a method involving a miniature microphone attached to the rat nasal cavity that allows to capture both male and female individual rat vocalizations and sniffing patterns during social interactions. We found that while the emission of 50 kHz USVs increases during close interactions, these signals lack specific behavioral associations. Moreover, a previously unreported low-frequency vocalization type marking rat social interactions was uncovered. Finally, different dynamics of sniffing and vocalization activities point to distinct underlying internal states. Thus, our method facilitates the exploration of internal states concurrent with social behaviors.

**Motivation:** Here, we sought to solve the problem of identifying the emitter of a specific vocalization during free social interaction of animal models. The use of a miniature microphone carried by the subject and connected to its nasal cavity solved this challenge and allowed for assignment of any vocalization recorded during a social interaction to a specific individual. Moreover, the miniature microphone enabled simultaneous recording of sniffing and vocalization activities of an individual, thus facilitating recognition of the internal states underlying these traits.

## Introduction

Auditory cues are a primary modality of social communication among animals, including humans ^1–3^. During social interactions, rodents emit mainly ultrasonic vocalizations (USVs) ^4^ that were shown to convey information regarding various environmental and internal conditions ^5,6^. Moreover, the rate and spectral characteristics of emitted USVs were found to be modified in various rodent models of neurodevelopmental disorders, such as autism spectrum disorder (ASD) ^7–9^. Adult laboratory rats primarily emit two types of USVs, prolonged 22 kHz calls, which reflect negative states, such as distress or alarm ^10,11^, and short 50 kHz calls that reflect positive states, such as affiliative social interactions or reward ^6,12^. Accordingly, playback experiments demonstrated that certain types of 50 kHz USVs induced approach behavior in adult rats, while 22kHz calls elicited avoidance responses ^13,14^. Thus, rat USVs can be used as a convenient way to assess the affective state of a subject under various conditions ^15^. Nevertheless, the use of USVs for studying the behavior and internal state of laboratory rodents have thus far been hindered by two main obstacles. First, USVs were traditionally detected and marked for further processing using manual analysis, which does not allow for high-throughput signal analysis. Second, it was almost impossible to separate calls from several animals grouped together using a standard microphone placed in the arena. While the first obstacle was recently solved using multiple computational tools that make use of machine learning algorithms to detect and analyze USVs (for comprehensive review see references ^16,17^), the second obstacle remained. Although recent studies using multiple arena microphones assigned USVs to specific individuals separated by a barrier ^18^, it remains almost impossible to distinguish and categorize USVs emitted by several individuals during free social interactions according to the emitter ^19–21^.

Rat breathing (or sniffing) is also indicative of the internal state of a subject, with the rate increasing from 1-4 Hz at rest to 8-12 Hz (theta range) during active exploration ^22^, reward anticipation ^23^ and arousal states ^24^. Specifically, the sniffing rate exhibits robust theta rhythmicity during social interactions ^18,25^. The most accurate experimental method for recording breathing in behaving rats involves a cannula implanted in the rat nasal cavity that is connected via a flexible tube to a pressure sensor located above the arena ^18,26^. Yet, the long elastic tube dampens the recorded pressure changes hence reduces their accuracy.

Here, we describe a method that allows simultaneous recording of both vocalization and sniffing activities of behaving individual animals using a miniature microphone carried by the subject animal. Using this method, we accurately detected and analyzed the vocalizations made by individual rats during free and restricted social interactions, even during close contact with other animals. We found that the tendency of the animals to vocalize was higher during close interactions, albeit without any association to a specific behavior. Moreover, we revealed a previously undescribed type of low-frequency weak vocalization emitted in parallel to 50 kHz USVs. We also measured and analyzed the sniffing rate and its relationship to the vocalization activity of a subject. Interestingly, the rates of USVs and sniffing produced by a subject rat presented different characteristic dynamics during social interactions, suggesting that distinct internal states drive these behavioral variables. The method presented here thus enables for the first time reliable relating of vocalizations to specific individuals and probing of the internal state of behaving animals during free social interactions by following both vocalization and sniffing activities. We hope that this method will enable for the monitoring of socio-emotional states in laboratory rats, including models of neurodevelopmental disorders, upon exposure to various environmental and internal conditions.

## Results

### Using a miniature microphone to record vocalizations of socially interacting individual rats

To separate ultrasonic vocalizations made by a subject rat from those made by its partner during social interaction, we attached a miniature ultrasonic microphone to the subject’s nasal cavity. To this end, we chronically implanted a cannula in the nasal cavity of the subject ^27^, and connected it by a short (3-5 cm) polyethylene tube to a miniature microphone placed on an Intan RHD2132 head-stage (Fig. 1A). The microphone signals were relayed to a data acquisition device and sampled at 250 kHz. Besides the miniature microphone used to monitor USVs made solely by the subject, a standard ultrasonic microphone placed above the arena (arena microphone) allowed us to record USVs produced by both animals (Fig. 1B).

**Figure 1.**
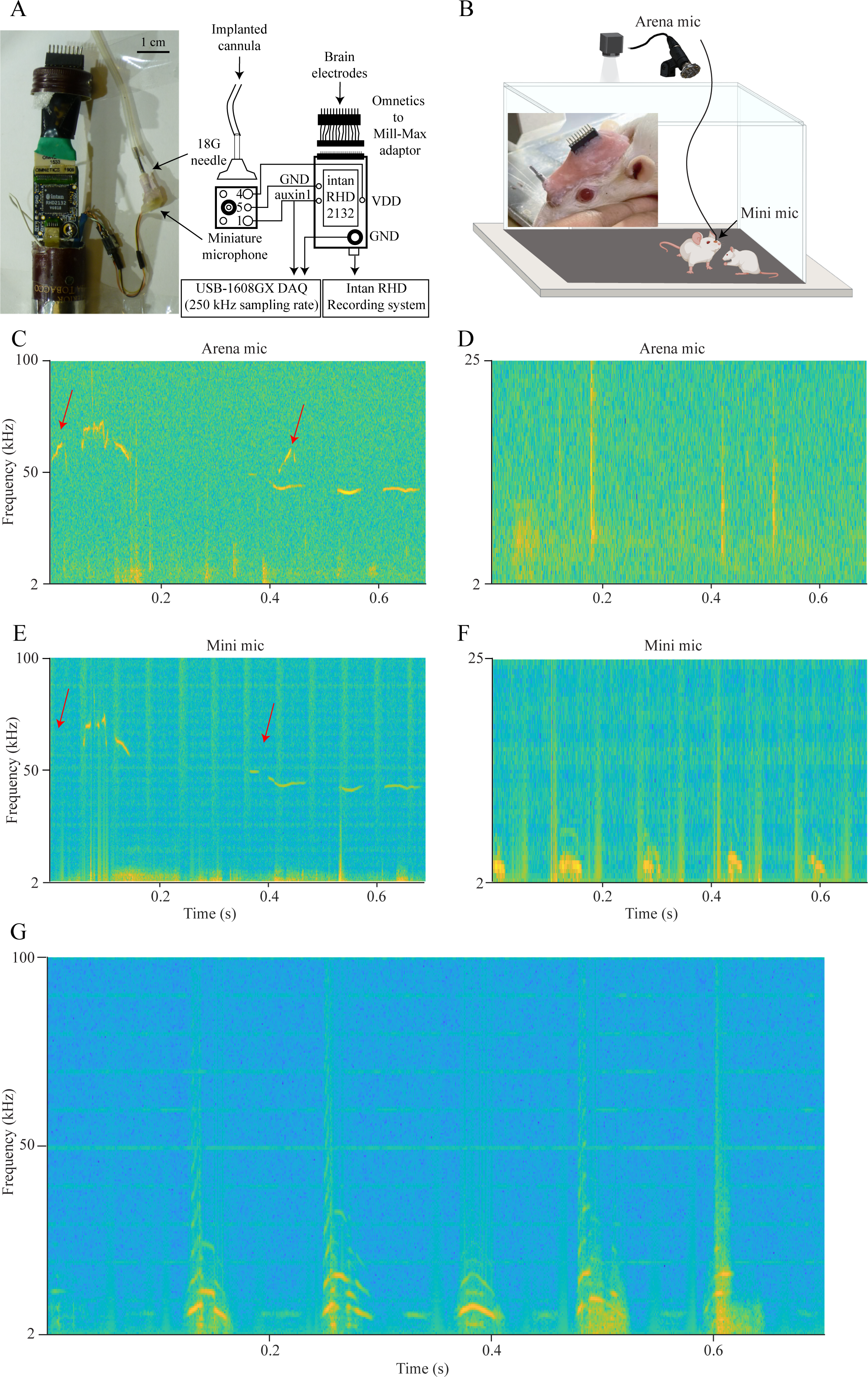
Recording rat social vocalization activity via a miniature microphone. **A.** A picture (left) and a scheme of the miniature microphone recording system, highlighting the various electronic and mechanical components. **B.** A schematic representation of the experimental setup showing the arena microphone located above the arena and the miniature microphone located on the subject’s head. The inset picture shows the rat head, including the connector and implanted cannula embedded in pink dental cement. **C.** A spectrogram showing various ultrasonic vocalizations made by both animals during a 0.7 s period of a social interaction, as recorded by the arena microphone. The USVs made by the stimulus animal are labeled by red arrows. **D.** A spectrogram of a different period with no USVs, as recorded by the arena microphone. **E.** The spectrogram of the same period shown in **C**, as recorded by the miniature microphone. Note that the stimulus calls, marked by red arrows, were not detected by the miniature microphone. **F.** The spectrogram of the same period shown in **D**, as recorded by the miniature microphone, which detected a burst of low-frequency vocalizations (LFVs). **G.** Another example of LFVs recorded using the miniature microphone, at higher resolution. Note the clear structure and harmonics of the LFVs. See also Fig. S5

We used this configuration to track social behavior and record USVs during 5 min of free social interactions (n=39 sessions) of adult male (n=8) and female (n=5) subject rats with either juvenile male or female partners (stimulus animals; Fig. S1A-E, Videos S1-2). We also recorded USVs during a social preference (SP) task ^28,29^ conducted by the same subject rats with either male or female stimulus animals and found normal preference of the animal over the object stimulus in all cases (n=32 sessions, Fig. S1F-J, Videos S3-4). As exemplified in Fig. 1C&E, USVs made by the subject could be detected by both microphones, while those made by the partner (marked by red arrows) were detected only by the arena microphone. Notably, the miniature microphone also recorded low-frequency (4-10 kHz) vocalizations (LFVs) that were not detected by the arena microphone (Fig. 1D, F), presumable because they were too weak. These previously unreported LFVs were typically emitted in regular bursts and had a clear structure in the frequency domain (Fig. 1G). Finally, in one free interaction session, we also encountered classical 22 kHz vocalizations, considered distress or alarm calls ^6,12^. These calls, emitted by a male stimulus animal in the presence of a female subject, were not further analyzed.

### The miniature microphone system can separate vocalizations of rats in close contact

Next, we validated that the separation of USVs according to the emitter was precise, even during close contact between the animals, a condition in which previous studies struggled to properly separate vocalizations made by freely interacting animals ^19–21^. For this, we analyzed all video files using TrackRodent software ^28^ and extracted all time epochs in which the subject animal either physically contacted the stimulus animal during free interaction (Fig. 2A) or investigated the chamber of the stimulus animal during a SP task (Fig. 2B). As shown in Fig. 2C-F, our system could reliably separate the USVs made by each animal, even during nose-to-nose contact (Fig. 2C, E) or upon investigation of the stimulus animal’s chamber by the subject (Fig. 2D, F). When analyzing the number of discrete vocalizations (syllables) across all sessions, we found a significant number of all three types of calls (i.e., subject USVs, stimulus USVs and subject LFVs) produced during physical contact in all types of experiments (Fig. 2G-L). Notably, the video and audio analyses were done in an automatic and unbiased manner. Thus, our system could adequately separate the various syllables according to their emitter in all examined conditions, regardless of the positions of the two animals. Moreover, when we analyzed recordings collected over a 5 min period after each session, when the stimulus animal was no longer in the arena ^30,31^, we could detect a substantial number of subject USVs and LFVs in most sessions (n=22 for USVs and 43 for LFVs), while encountering only one USV mistakenly annotated as a stimulus USV (Fig. 2M). This mistake was most probably due to temporary blockage of the cannula, which caused this subject USV not to be detected by the miniature microphone, and hence annotated as a stimulus USV. Thus, the error rate of our annotation was <1% (1 out of 213 USVs).

**Figure 2.**
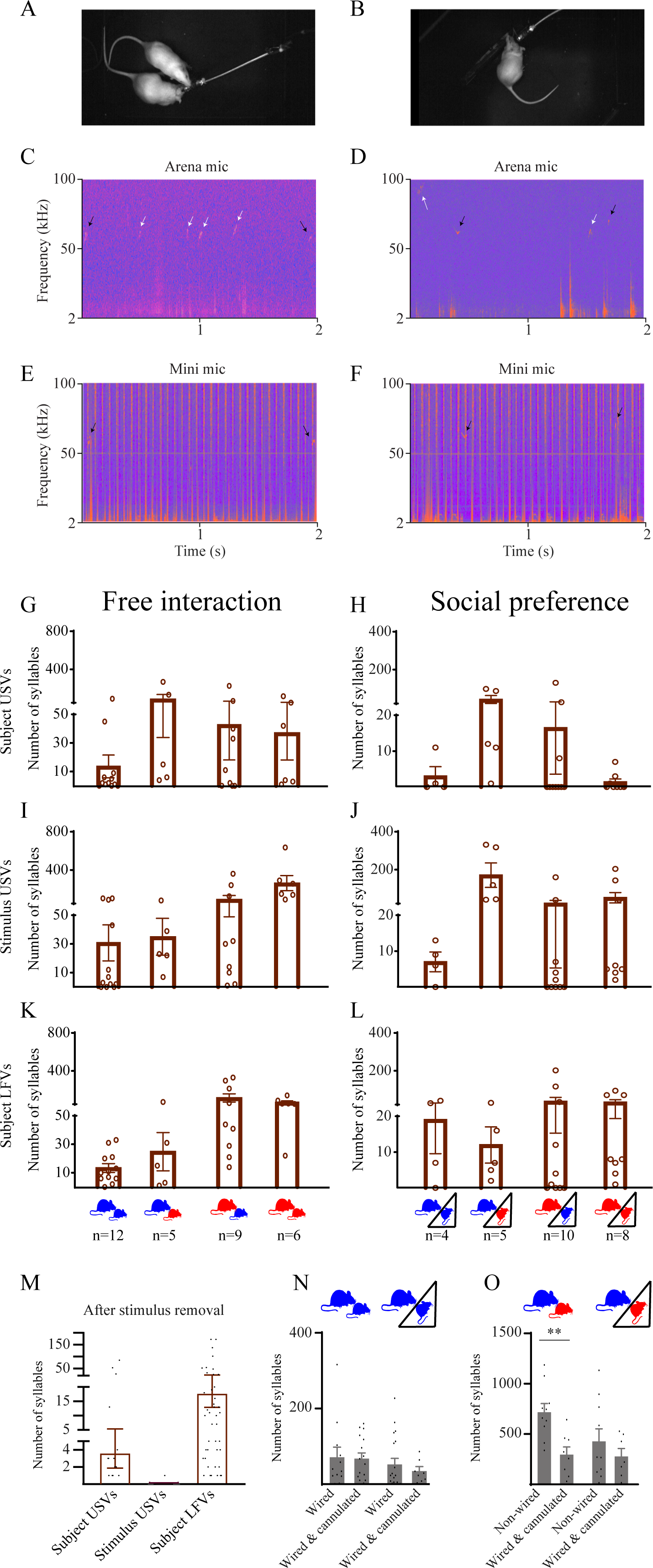
Identification of social call emitted by individual rats in close contact. **A.** A picture of two rats in close nose-to-nose contact during free social interaction. Note the bright cable extended from the subject’s head. **B.** A picture of two rats in close nose-to-nose contact during a social preference task. **C.** A 2 s spectrogram of vocalizations detected by the arena microphone during the free interaction depicted in **A**. White arrows label USVs emitted by the stimulus animal while black arrows point to vocalizations emitted by the subject, also detected by the miniature microphone (see **E**). **D.** As in **C**, for the SP interaction shown in **B**. **E.** The spectrogram of the miniature microphone during the same time as in **A** and **C**. Note that only the subject USVs labeled by black arrows are detected. **F.** As in **E**, for the event shown in **B** and **D**. **G.** Mean (±SEM) number of subject USVs detected during physical contact between the subject and stimulus animals during free social interaction with various male and female combinations (see below (**K**) for X-axis labels: blue-male, red–female, big–subject, small–stimulus animal). **H.** As in **G**, for SP sessions (see below (**L**) for definition of male/female combinations). **I.** As in **G**, for stimulus USVs. **J.** As in **I,** for SP sessions. **K.** As in **G**, for LFVs. The depictions below show male/female combinations and the number of sessions for each bar (blue-male, red–female, big–subject, small– stimulus animal). **L.** As in K, for SP sessions. **M.** Mean (±SEM) number of the various calls detected during the 5 min period following stimulus removal from the arena after both free interaction and SP sessions, when subject animals are still looking for the stimulus animal, which is no longer present, and hence, cannot vocalize. Note that only one USV (out of 213) was mistakenly annotated as a stimulus USV during this period. **N.** Mean (±SEM) number of USVs detected by the arena microphone during either free interaction (left two bars) or SP (right two bars) male-male interactions, with subject that were either just wired with a cable to the recording system or wired and implanted with a cannula in their noses. **O.** As in **N**, for interactions between male subjects and female stimulus animals (see depictions above, blue-male, red–female, big–subject, small–stimulus animal), with the subjects being either non-wired (freely moving) or wired and implanted with a cannula. **p<0.01, Student’s t-test. For detailed statistical analysis results, see Table S1.

Finally, to assess if the implanted cannula caused any reduction in USV production, we compared the number of USVs detected by the arena microphone during either free interaction or a SP task conducted by male subjects and stimulus animals, with the subject either being cannulated and wired or just wired to the recording system. We found no significant difference in the number of USVs emitted by cannulated vs. wired animals, suggesting that the cannula itself did not interfere with vocalization activity (Fig. 2N; p>0.2, Student’s t-test). However, when comparing the number of USVs produced by wired and cannulated vs. non-cannulated freely moving males during interaction with female stimulus animals, we found a significant reduction in the case of free interaction (t_16_ = 3.668, p<0.01) but not during the SP task (Fig. 2O; t_15_ = 0.9230, p>0.2, Student’s t-test). Thus, it seems as if wired animals indeed produce less calls than do freely moving animals in certain contexts.

Overall, these results suggest that our method enables accurate detection and separation of vocalizations according to their emitter during social interaction in various conditions and contexts, regardless of the animals’ distance from each other.

### Analyzing the number and characteristics of social vocalizations in various experimental and social contexts

We then compared the number of syllables made either by the subject or stimulus animal during free interaction sessions in all subject/stimulus combinations (social contexts, Fig. 3A-D). While female subjects made significantly more physical contacts with either male or female stimulus animals than did male subjects (Fig. 3A), we found that male, but not female subjects, emitted significantly more USVs during interactions with female than with male stimulus animals (Fig. 3B). In parallel, Female stimulus animals emitted significantly more USVs in the presence of a female subject compared to that in the presence of a male subject, and to male stimulus animals in the presence of a female subject (Fig. 3C). Finally, we found that female subjects consistently made more LFVs than male subjects when interacting with male stimulus animals (Fig. 3D).

**Figure 3.**
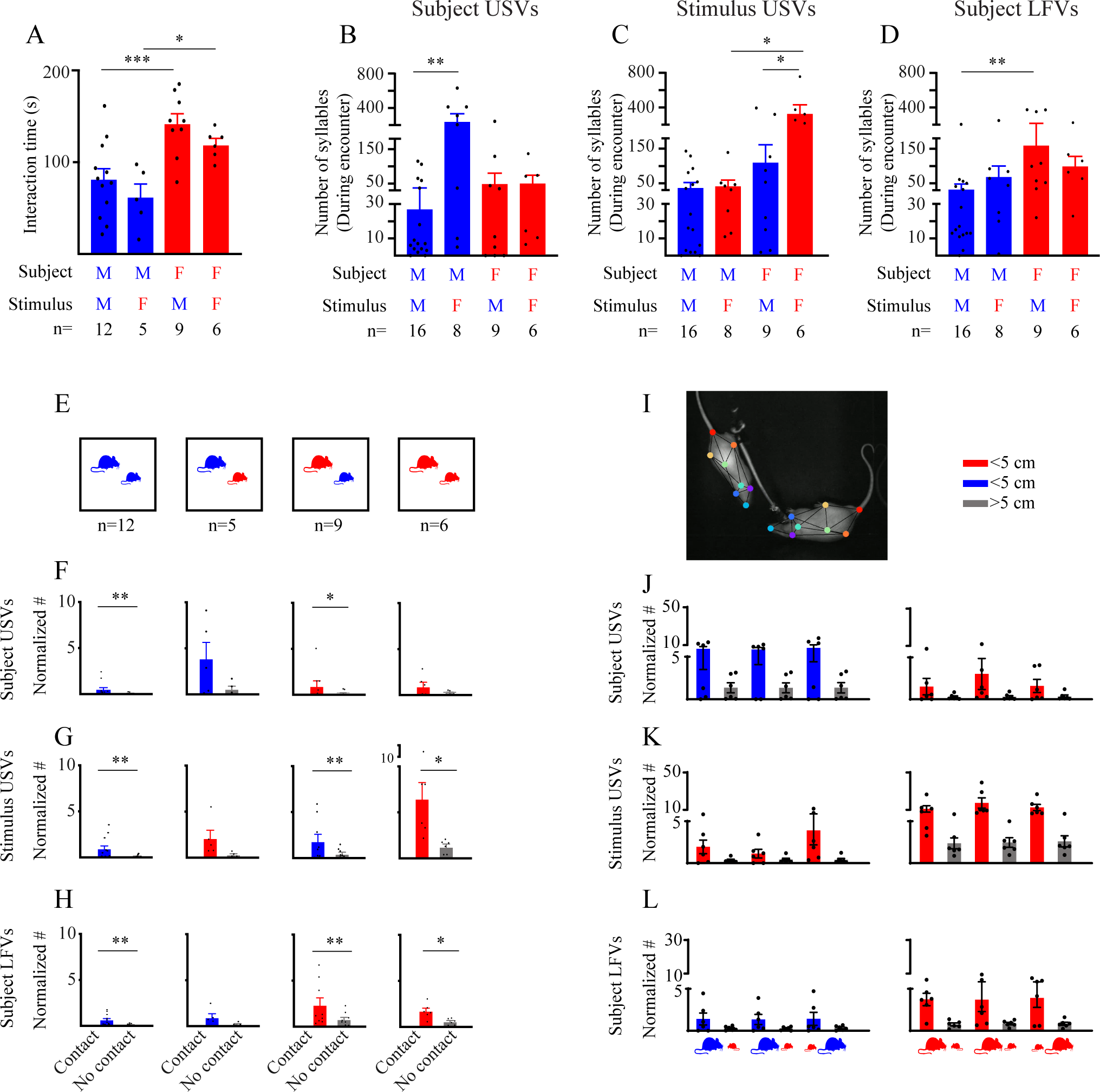
Vocalization rates are higher during close interactions but are not linked to a specific behavior. **A.** Mean (±SEM) interaction (physical contact) time during free social interactions of the various male/female combinations (social contexts). The sex of subject and stimulus animal in each context (blue-male, red–female, big–subject, small– stimulus animal), as well as the number of animals (n) appear below. *p<0.05, **p<0.01, ***p<0.001, *post hoc* Sidak’s multiple comparison test following significant main effect in a two-way ANOVA test. **B.** As in **A**, for the number of subject USVs. **C.** As in **A**, for stimulus USVs. **D.** As in **A**, for stimulus LFVs. **E.** Schemes of the various social contexts of the free social interaction sessions (blue-male, red–female, big–subject, small–stimulus animal). The number of sessions for each context is noted below. **F.** Mean (±SEM) number of subject USVs emitted while the animals were in physical contact (left bar) or without physical contact (right bar), normalized to the time spent in contact or with no contact, in each social context. *p<0.05, **p<0.01, paired t-test. **G.** As in **F**, for stimulus USVs. **H.** As in **F**, for subject LFVs. **I.** An example picture of DeepLabCut pose estimation during a free social interaction session. **J.** Mean (±SEM) number of subject USVs (normalized as in **F**) emitted while the animals were in <5 cm (blue-male, red-female) or >5 cm (grey) nose-to-nose (left two bars), nose-to-tail (middle two bras), or tail-to-nose (right two bars, see depictions below **L,** blue-male, red–female, big–subject, small–stimulus animal) position. Two-way ANOVA, main effect of distance, p<0.01 and p<0.05 for males and females, respectively. **K.** As in **J**, for stimulus USVs. Two-way ANOVA, main effect of distance, p<0.01 and p<0.05 for males and females, respectively. **L.** As in **J**, for subject LFVs. Two-way ANOVA, main effect of distance, p<0.05 and p<0.0001 for males and females, respectively. For detailed statistical analysis results, see Table S1. See also Figs. S1, S2 and S4.

Overall, these results suggest that vocalizations of all types are emitted during free social interaction in a sex- and social context-dependent manner by both partners. Similar conclusions can be drawn from the same analysis conducted during SP sessions (Fig. S2). Moreover, we observed significant sex- and social context-dependent changes in some of the spectral characteristics of the various calls, such as their mean frequency and duration (Fig. S3).

### Social vocalizations are more common during close interaction, irrespective of specific behavior

Social vocalizations can be associated with close interaction between animals or may merely reflect social context. In the latter case, they are expected to appear equally during periods of contact vs. no contact between the animals. We, therefore, normalized the number of vocalizations emitted in free interaction sessions while the animals were either in physical contact or without contact, by dividing these numbers by the time spent by the animals either in contact or without contact, respectively. We found that in most cases, there was a significantly higher normalized number of both subject and stimulus USVs during physical contact than during no-contact periods. This suggests that at least part of the USVs were associated with the social interaction itself (Fig. 3E-H). We thus concluded that rats show a higher tendency to emit vocalizations during close social interaction and that these calls are not just a reflection of the social context.

Still, USVs may be associated with specific behavioral events that occur during social interaction or are emitted with no correlation to a specific behavioral event. To distinguish between these possibility, we used DeepLabCut ^32,33^ (Fig. 3I) to analyze the distance between animals during free interaction in contexts characterized by many vocalization, in three distinct behavioral events, namely, nose-to-nose, nose-to-tail (subject following the stimulus) and tail-to-nose (subject followed by the stimulus) interactions. Here too, we normalized the number of vocalizations emitted while the animals were in each type of behavioral event (Fig. S4A-C), by dividing it with the time spent by the animals in the respective behavioral situation (Fig. S4D). In agreement with the results obtained using TrackRodent (Fig 3E-H), we found that all three types of calls had significantly higher probability of appearing during close (<5 cm) interaction than during remote interaction (Fig. 3J-L; p<0.05, two-way ANOVA for each context). However, there were no differences among the three types of behavioral events (p>0.2, two-way ANOVA for each context). Thus, it seems as if rat social vocalizations are part of their social interactions but are not associated with a specific behavioral event.

### The miniature microphone enables highly sensitive recording of sniffing activity

The miniature microphone recorded all pressure changes in a subject’s nasal cavity, thus allowing us to monitor not only vocalizations but also the sniffing activity of the subject during the recorded sessions. To extract the sniffing signal, we relayed the miniature microphone output to the Intan head-stage (AUX input) and analyzed these signals after down-sampling them at 1 kHz (Videos S1-4). In agreement with previous studies ^18^, we observed a relatively low rate (4-7 Hz) and amplitude of sniffing during the baseline period (Fig. 4A). In contrast, a high sniffing rate (10-15 Hz) and amplitude were observed throughout the social interaction (Fig. 4B), although a moderate reduction occurred towards the end of the encounter (Fig. 4C-D). Notably, the sniffing rate and amplitude started to increase several tens of seconds before stimulus introduction (Fig. 4C), most probably triggered by detection of the experimenter’s preparations (about 30 s before stimulus introduction time).

**Figure 4.**
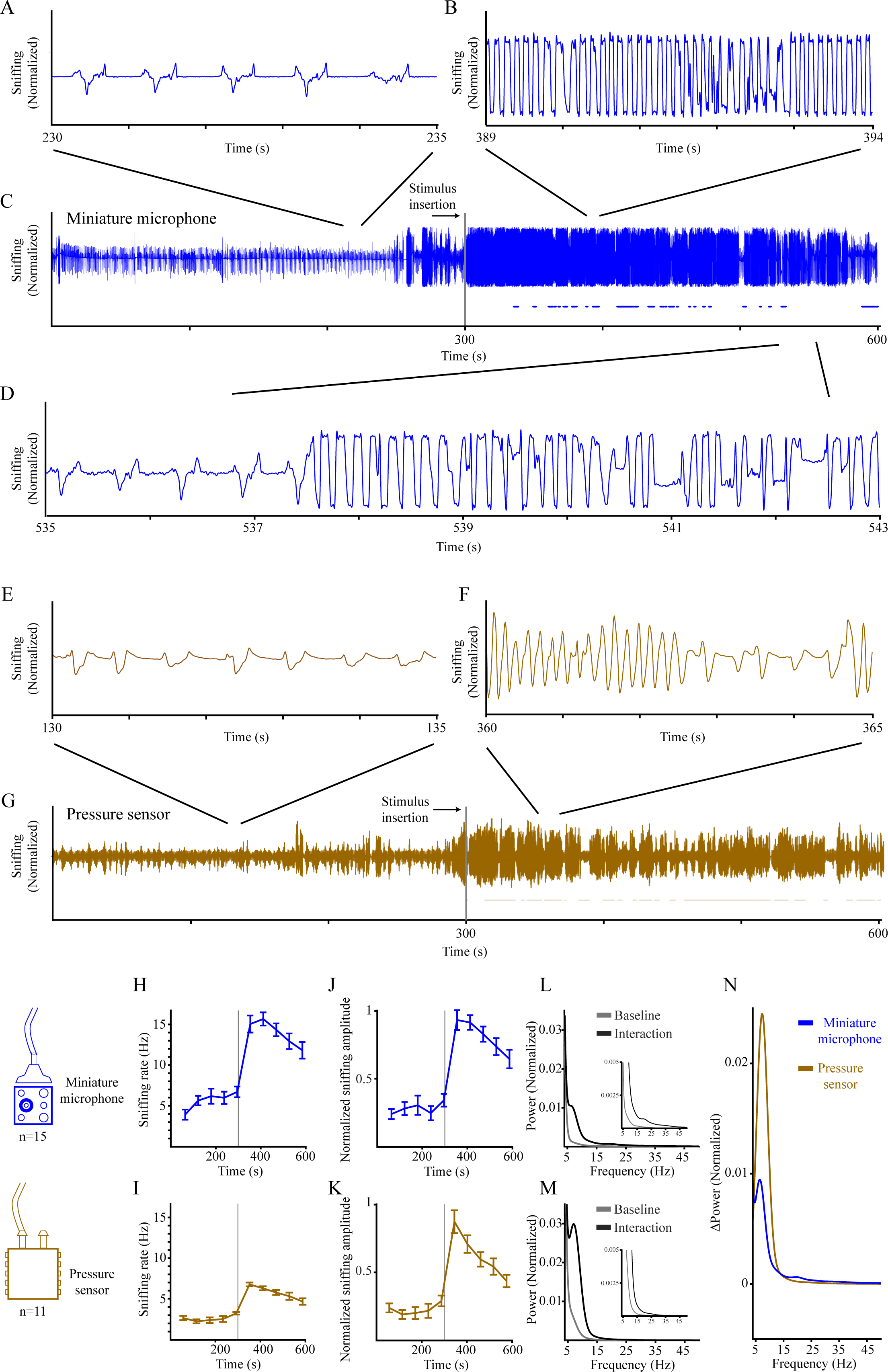
Recording sniffing activity using the miniature microphone. **A.** A 5 s trace of pressure changes in the nasal cavity (sniffing activity) recorded from a male rat using the miniature microphone during baseline, about 70 s before stimulus introduction into the arena. Note the low amplitude and rate of the sniffing activity at this stage of the session. **B.** As in **A**, for sniffing activity of the same animal recorded about 90 s after the introduction of the stimulus animal into the arena. Note the high rate and amplitude of the sniffing activity at this stage of the experiment. Also, note the trimming of activity peaks and troughs by the system, due to the strong signal. **C.** Recording of the whole session, including the periods shown in **A** and **B**. Note the decline in sniffing rate and amplitude about 200 s after stimulus introduction. **D.** An 8 s trace taken from the same session, towards the end of the encounter, showing intermittent periods of low and high sniffing rates. **E.** As in **A**, for the signal recorded using a pressure sensor from a different animal, about 150 s before stimulus introduction. **F.** As in **E**, recorded about 60 s following introduction of the stimulus animal. Note that due to the weaker signal of the pressure sensor, it was not trimmed by the recording system. **G.** Recording of the whole session, including the periods shown in **E** and **F**, using the pressure sensor. **H.** Mean (±SEM) sniffing rate analyzed over the course of the whole session (baseline and interaction) using 1 min time bins, as recorded from eight males using the miniature microphone. **I.** As in **H**, for four males recorded using the pressure sensor. Note the lower range but similar dynamics, as compared to **H**. **J.** As in **H**, for normalized sniffing amplitude. **K.** As in **I**, for normalized sniffing amplitude. **L.** Superimposed power spectral density (PSD) profiles for signals recorded using the miniature microphone during the baseline (grey) and interaction (black) periods. The inset shows PSD profiles at higher power resolution. **M.** As in **L**, for signals recorded using pressure sensor. **N.** Superimposed PSD profiles of the miniature microphone (blue) and pressure sensor (gold) signals, after subtraction of the baseline from the encounter, separately for each signal. Note the higher power recorded by the pressure sensor in the theta (4-12 Hz) range, as compared to the higher power recorded by the miniature microphone in the 15-25 Hz range. For detailed statistical analysis results, see Table S1. See also Fig. S5.

To directly compare pressure recordings made by the miniature microphone with the more commonly used pressure sensor ^18,26,27^, we conducted several experiments (n=4 male subjects, 11 sessions) with a pressure sensor coupled to the implanted cannula (Fig. S5). This yielded qualitatively similar signals to those detected by the miniature microphone (Fig. 4E-G). Notably, the miniature microphone yielded stronger signals that surpassed the range of the Intan system’s AUX input. Hence, the system trimmed its peaks and troughs. Nevertheless, at midrange, pressure changes were recorded at higher resolution by the miniature microphone, as compared to the pressure sensor (compare traces of both techniques in Figs. 4B and F). Although we observed very similar dynamics of sniffing rate and amplitude using both devices (Fig. 4H-K), the sniffing rate calculated from the peaks of the pressure changes detected by the miniature microphone (Fig. 4H) was higher than that calculated from the pressure sensor signal, which ranged from 2-4 Hz at baseline to 6-8 Hz during the encounter (Fig. 4I). This difference was most probably due to the higher resolution of the sniffing signal detected by the miniature microphone, which allowed detection of local peaks that were overlooked by the pressure sensor. To confirm that the difference between the two techniques was not due to vocalizations recorded by the miniature microphone, we calculated the normalized power spectral density (PSD) profiles of the signals before (baseline) and during interaction, after removing all time segments that contained USVs. The results clearly showed how both techniques yielded increased PSD in the 4-50 Hz range during the encounter, as compared to the baseline period (Fig. 4L-M). However, the change in theta-range power (4-12 Hz) was more prominent in the pressure sensor signal than in the signal recorded by the miniature microphone, whereas the PSD of the miniature microphone was more robust at higher frequencies (15-40 Hz) (Fig. 4N). These results suggest that the miniature microphone detected subtle high-frequency pressure changes better than did the pressure sensor.

### Distinct dynamics of vocalization and sniffing activities during social interaction

Finally, since both the 50 kHz USVs and high sniffing rate are thought to be associated with internal states, such as attention and arousal ^34–36^, we assessed whether sniffing and vocalization activities showed similar dynamics along all sessions. Were similar dynamics to be observed, this would suggest that a single state drives them all. As shown in Fig. 5A, the rate of subject USVs started to rise only after introduction of the stimulus animal. Accordingly, there was no significant difference in the number of USVs recorded during a 20 s interval before stimulus introduction and the mean number of USVs recorded during the earlier 280 s baseline period (Fig. 5B). In contrast, the number of LFVs clearly started to rise before stimulus introduction (Fig. 5C) and the number of vocalizations during the 20 s period before stimulus introduction was significantly higher than the mean number during the baseline period (Fig. 5D). Similarly, both the rate and amplitude of sniffing activity started to rise before stimulus introduction (Fig. 5E-H). However, while the sniffing variable significantly declined over the course of the session (Fig. 5I-J), both USVs and LFVs did not show such decline (Fig. 5K-L). In fact, the USV rate showed a tendency to increase along the session, although this increase did not reach statistical significance in a non-parametric test. We, therefore, concluded that sniffing activity and the two types of social vocalizations were being driven by distinct internal states. Notably, we did not find any difference in sniffing dynamics between the first session experienced by the subject and subsequent sessions (Fig. S6), suggesting that it is not the expectation of a social encounter that drives the elevation in sniffing activity, but rather a distinct state, such as attention, which is triggered by the experimenter’s preparations.

**Figure 5.**
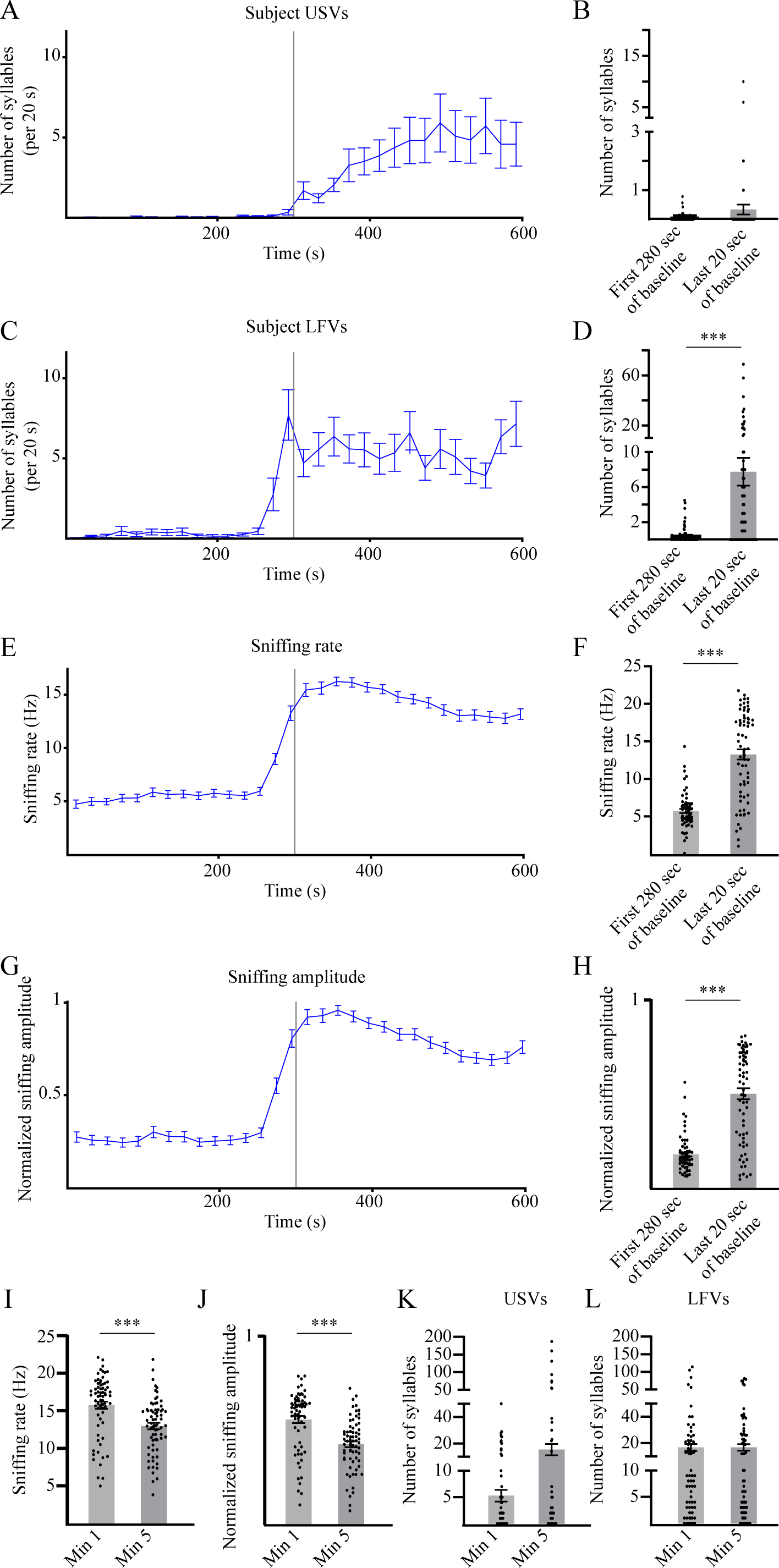
Distinct dynamics of vocalization and sniffing activities during social interaction. **A.** Mean (±SEM) number of subject USVs (20 s bins) produced before (left to the vertical line) and after (right to the vertical line) introduction of the stimulus animal. **B.** Mean (±SEM) total number of subject USVs (20 s bins) produced during the 280-s period from the beginning of the recording till 20 before stimulus introduction (left bar) and during the last 20 s before stimulus introduction (right bar). **C-D**. As in **A-B**, for subject LFVs. Note the significant increase in LFVs production occurring 20 s before stimulus introduction. **E-F**. As in **C-D**, for subject sniffing rate. **G-H**. As in **C-D**, for subject sniffing amplitude (normalized). **I.** Mean (±SEM) sniffing rate during the 1st (left bar) and 5^th^ (right bar) minute after stimulus introduction. Note the significant difference between the two periods, suggesting a decrease in sniffing rate along the social interaction. **J.** As in **I**, for the normalized sniffing amplitude. **K.** As in **I**, for the number of subject USVs. Note the lack of reduction along the social interaction. **L.** As in **K**, for the number of subject LFVs. For detailed statistical analysis results, see Table S1. See also Fig. S6.

Overall, these results validate the miniature microphone as a highly sensitive means for monitoring vocalization and sniffing activities from individual rats during social interactions.

## Discussion

Here, we present a method that allows simultaneous recordings of sniffing and vocalization activities from individual animals during various types of social interactions. Since this method is based on a miniature microphone directly connected to the subject’s nasal cavity, it can be reliably used to detect the vocalization and sniffing activities of the subject even during free social interaction, regardless of the subject’s position relative to other animals. We also demonstrated that recording from an arena microphone in parallel to the miniature microphone allowed precise separation of the vocalizations of two animals during free social interaction.

Previous studies employed multiple indirect ways to separate the vocalizations of two animals during social interaction. In some cases, the animals were separated into distinct compartments by a barrier, and recordings were conducted using two microphones, each located in a different compartment, thus allowing separation of vocalizations according to the emitter ^18,37^. This arrangement, however, restricts the types of interaction between animals and does not allow them to freely contact each other. Other studies combined an array of microphones around the arena and performed computational analysis of the various signals to decide which animal emitted a given USV ^19–21^. The efficiency of this technique was limited to cases where the heads of the animals were distant from each other by several centimeters (up to ∼1.5 cm with accuracy of 84.3% ^21^). As far as we know, our system is the first one that allows for fully reliable direct recording of vocalization made by each animal separately, regardless of its position relative to other animals.

In general, rat 50 kHz USVs are associated with positive states and rewarding contexts, such as affiliative social behaviors ^6,14,38^. The role of such USVs, however, is not yet clear. They could simply reflect the subject’s internal state, with no relation to immediate interactions, be associated with the interaction or be directly linked to a specific behavioral event, such as social investigation or following. Distinguish these possibilities would be extremely difficult using previously published methods, as they do not allow for annotation of the USVs emitted by a specific individual during free social interaction. We thus employed our method to address this question and found that a higher tendency of both subject and stimulus animals to vocalize while close to each other during both free social interaction and a SP task. However, this was just a tendency, as the animals were vocalizing even when they were remote from each other. We used DeepLabCut to assess whether the tendency for USV production was higher during relevant behavioral events, such as nose-to-nose or nose-to-tail interactions, and found no such link. We, therefore, conclude that rat tendency to emit 50 kHz USVs in social contexts is higher during instantaneous interactions, yet is not linked to any specific behavioral event we could recognize. These results, which are in agreement with previous studies using devocalization experiments ^39–40^, suggest that rat 50 kHz USVs reflect a positive affective state that is enhanced when the animals are in close interaction and are not part of a specific type of behavior.

We further used our system to analyze the number of USVs in various combinations of males and females and found that while male subjects vocalized significantly more towards female stimulus animals, female subjects showed similar vocalizations towards both male and female stimulus animals. In contrast, female stimulus animals emitted significantly more vocalizations while interacting with female than with male subjects. We also found small but statistically significant differences in the spectral characteristics of the recorded vocalizations across the various social contexts. Overall, these results add to multiple recent studies demonstrating that rat USVs are age-sex- and social status-dependent, and hence may convey social information during social interactions ^41–45^.

Using the high sensitivity of the miniature microphone recordings, we revealed a previously unreported type of low-frequency vocalization (LFV) that had not been reported by previous studies. These vocalizations seem to be too weak to be detected by arena microphones, which may explain why they were not reported so far. Similar to USVs, the LFVs were detected almost only during social interactions and exhibited dependencies in terms of rate and spectral features on the social context (sex combination) of the interaction, suggesting that these vocalizations are also involved in social behavior. Notably, LFVs were detected in the same exact recordings as USVs but did not overlap with the USVs and had a very different frequency range (4-10 kHz). Moreover, the LFVs presented distinct dynamics from USVs, as the production of the former started to rise even before introducing the stimulus into the arena, as also seen with sniffing activity, whereas the production of subject USVs started only after stimulus introduction. Thus, LFVs cannot be an artifact of USVs production. Future studies may reveal whether LFVs play a specific role during social interactions. It should be noted that somewhat similar “mid-frequency” vocalizations were previously reported in mice following restraining ^46^. However, in our experiments, we hardly detected any distress 22 kHz calls, with the subject rats showing robust social preference in the SP task. Thus, it is most likely that rat LFVs are part of affiliative social interactions, similar to 50 kHz USVs. Nevertheless, the different dynamics of LFVs and USVs suggests that they reflect distinct internal states associated with affiliative social interactions.

We have also demonstrated that the miniature microphone enabled reliable measurements of pressure changes in the subject’s nasal cavity at a higher resolution than the previously used pressure sensors ^18^. Exploiting this ability revealed that sniffing and vocalization activities followed distinct dynamics. Whereas sniffing rate and amplitude started to rise as early as several tens of seconds before stimulus introduction, the number of 50 kHz USVs started to increase only after stimulus introduction. Moreover, while sniffing activity peaked within one minute following stimulus introduction and then gradually decreased, vocalization activity of both the subject and the stimulus did not decrease and even increased over the session. Thus, while both activities were previously suggested to reflect an internal state of arousal ^35,47^, our results suggest that in the context of social encounters between adult and juvenile rats, they reflect distinct internal states. We propose that sniffing activity is mainly modulated by attention, with its rate starting to rise as soon as the animal detects an environmental change, such as the preparations of the experimenters to introduce stimuli into the arena. This proposal is in accordance with previous studies demonstrating higher sniffing rates in association with states of attention and exploration in adult rats ^48–51^. In contrast, 50 kHz USVs seem to reflect social bonding processes ^35^, and thus start only after introducing the stimulus animal into the arena. In some cases, these USVs gradually increased (see Fig. S1). Notably, unlike previous studies showing vocalization activity due to reward anticipation ^12,52–39^, we did not observe significant amount of 50 kHz USVs before the introduction of the stimulus animal, even with subjects that had already experienced sessions of social interaction in the arena. This discrepancy may be explained by strain-, age-, context- and experience-dependent differences among the various studies.

Overall, our method of recording rat vocalization and sniffing activities using an implanted miniature microphone generated valid and accurate data which allow for exploration of both activities from individual rats during free social interaction at a higher resolution than attained with previous techniques. Therefore, this method should facilitate the use of these behavioral variables for characterizing social interactions in various animal models and conditions.

### Limitations of the study

One clear limitation of the study is that it involves surgery, cannula implantation and connecting of the subject to a recording system via electrical cables, all of which can reduce the tendency of the animal to produce USVs. While we found that cannula implantation itself did not cause a reduction in USV production, as compared to animals who were wired to a recording system without the cannula, we also noted that wired animals produced less USVs than did freely moving animals conducting the same task. This limitation may be solved by the use of wireless recording systems. Moreover, in combination with wireless recording systems that would overcome the problem of multiple wires in the arena, our method could allow for future recordings of sniffing and vocalization activities from groups of freely behaving animals, which is almost impossible to achieve using arena microphones.

Another limitation of our method is that it may be less efficient for recording small animals, such as mice or juvenile rats. We have tried using our approach with adult mice and found that the cannula rapidly clogged, and thus could not be used for extended recordings as with adult rats. Future improvements to our method may overcome this issue.

An advantage and potential application of our method is related to its enabling efficient combination of audio and sniffing recordings with invasive exploration of brain activity, as our setup is interfaced with the Intan RHD 2000 electrophysiological recording system. This will allow for recording of brain activity during various tasks while continuously assessing the subject’s internal state as reflected by its vocalization and sniffing activities.

## Declarations

### Acknowledgments

This study was supported by the ISF-NSFC joint research program (grant No. 3459/20 to S.W.), the Israel Science Foundation (grants No. 1361/17 and 2220/22 to S.W.), the Ministry of Science, Technology and Space of Israel (Grant No. 3-12068 to S.W.), the DFG (GR 3619/16-1 and SH 752/2-1 to S.W.) and the United States-Israel Binational Science Foundation (grant No. 2019186 to S.W.).

### Author contribution

Conceptualization, S.N. and S.W.; Methodology, A.B. and S.N.; Software, Y.G. and S.N.; Formal Analysis, R.T. and S.N., Investigation, R.A., R.T., S.R.J., Resources, A.B. and S.N.; Data Curation; Writing, S.N. and S.W.; Visualization, S.N.; Supervision, S.N. and S.W.; Project administration, S.N.; Funding Acquisition, S.W.

### Competing inter1ests

The authors declare no competing interests.

### Inclusion and diversity

We support inclusive, diverse, and equitable conduct of research

### Data availability

All processed data of the various recordings appear in Table S2.

## STAR METHODS

### RESOURCE AVAILABILITY

#### Lead Contact

Further information and requests for resources and reagents should be directed to and will be fulfilled by the lead contact, Shai Netser.

#### Materials availability

This study did not generate any new or unique reagents.

#### Data and Code availability

- The processed datasets analyzed during the current study (Table S2), as well as the detailed statistical analyses (Table S1), are deposited at Mendeley Data using the following reference: John, shanah; Tiwari, Rishika; Goussha, Yizahq; Amar, Rotem; Bizer, Alex; Netser, Shai; Wagner, Shlomo (2023), “Results and statistical summary for the paper - Simultaneous recording of ultrasonic vocalizations and sniffing from socially interacting individual rats using a miniature microphone”, Mendeley Data, V1, doi: 10.17632/whmf6d7cv7.1
- All original code has been deposited at Zenodo and Github and is publicly available as of the date of publication. DOIs are listed in the key resources table.
- Any additional information required to reanalyze the data reported in this paper is available from the lead contact upon request.

## EXPERIMENTAL MODEL AND SUBJECT DETAILS

### Animals

All animals were kept in the rat facility of the University of Haifa under veterinary supervision, in a 12 h light/12 h dark cycle (lights on at 9 PM), with *ad libitum* access to food (standard chow diet, Envigo RMS, Israel) and water. Subjects were Sprague Dawley (SD) male (n=12, eight recorded with a miniature microphone and four with a pressure sensor) and female (n=5) rats (9-15 weeks old) grown in-house and kept in groups of 2-5 animals per cage, until surgery. Rat stimulus animals were in-house-grown SD juvenile male and female rats (5-8 weeks old), kept in groups of 2-5 animals per cage throughout the experiment. Before cannula implantation surgery, subject rats were handled daily for 1-2 weeks. After implantation, the animals were isolated for about seven days throughout the following week of experiments. Behavioral experiments occurred during the dark phase of the light/dark cycle under dim red light.

### Institutional review board

All experiments were performed according to the National Institutes of Health guide for the care and use of laboratory animals and approved by the Institutional Animal Care and Use Committee (IACUC) of the University of Haifa.

## METHOD DETAILS

### Experimental setups

The experimental setup was as previously described ^28^. Briefly, a black matte Plexiglas arena (50 X 50 X 40 cm) was placed in the middle of an acoustic chamber (90 X 60 X 85 cm), which was electrically shielded and grounded to the recording systems using 2 mm aluminum plates. A high-quality monochromatic camera (Flea3 USB3, Point Grey), equipped with a wide-angle lens (Fujinon 6 mm fixed focal length C-mount lens, Point Grey), was placed at the top of the acoustic chamber and connected to a computer, enabling a clear view and recording (30 frames/s) of subject behavior using commercial software (FlyCapture2, Point Grey). For SP experiments, two black Plexiglas triangular chambers (20.5 cm isosceles, 40 cm height) were placed in two randomly selected corners of the arena, with a metal mesh (25 X 7 cm, 2.5 X 1 cm holes) covering the bottom of the triangular chamber.

### Behavioral paradigm

Each subject animal was tested twice a day for five consecutive days, with either free interaction or a SP task using either male or female stimulus animals, in random order. The stimulus animals were always novel to the subjects at the beginning of the experiment.

#### Free interaction

The behavioral paradigm started with a 15 min habituation of the subject rat to the arena. Next, the recording started for an additional 5 minutes of baseline recording, followed by a 5 min test of free interaction with a novel stimulus animal (see Fig. 1B for schematic description). Following the task, the stimulus was removed from the arena, and the subject rat was recorded in the arena for an additional 5 min period.

#### SP task

The experiment started with a 15 min habituation to the arena containing two empty chambers. Throughout this time, stimulus animals were placed in other chambers for acclimation. The recording session started with an additional 5 min of baseline with the empty chambers. Thereafter, the empty chambers were replaced with the animal and object (plastic toy, ∼5×5 cm) stimulus chambers, and the SP task was performed for 5 min. Following the task, the chambers with the stimuli were replaced with empty chambers, and the subject rat was recorded in the arena for an additional 5 min period.

### Surgery

Rat subjects were anesthetized by intraperitoneal injection of a mixture of ketamine and Domitor (0.09 mg/gr and 0.0055 mg/gr, respectively), or isoflurane through a low-flow anesthesia system (0.5–1%, ∼200 ml/min; SomnoFlo, Kent Scientific) and the painkiller Norocarp (0.016 mg/g). The level of anesthesia was monitored by testing toe pinch reflexes. The animals’ body temperature was kept constant at approximately 37°C using a closed-loop custom-made temperature controller connected to a temperature probe and a heating pad placed under the animal. Anesthetized animals were fixed in a stereotaxic apparatus (Stoelting Inst.), with the head flat. The skin was shaved and then gently removed, and a holes were slowly drilled into the nasal cavity to implant the cannula (16 G blunt needle, 11 mm length) and screws. The implanted cannula and screws were fixed with dental cement, in which we also dipped a Mill-Max connector (853-43-100-10-001000, Mill-Max) for holding the Intan recording head-stage. Immediately after surgery, the open cannula was tested by connecting it to a pressure sensor (MB-LPS1-01-100U5N, Microbridge) connected to an oscilloscope (ADS1013D, DANIU), to make sure the cannula was not clogged. After the test, the cannula was filled with a dummy blunt needle (20 G blunt needle, 12-12.5 mm length) to keep it clean. Following surgery, animals received daily injections of Norocarp (0.016 mg/g) for three days and were allowed to recover for at least five days before experiments.

Animals without implanted cannula that were tested while wired were treated as previously described ^55^.

### Audio recordings

#### Arena microphone

Vocalizations were recorded using a condenser ultrasound microphone (CM16/CMPA, Avisoft) placed high enough (∼80 cm) above the experimental arena such that the receiving angle of the microphone covered the whole arena. The microphone was connected to an ultra-sound recording interface (UltraSoundGate 116Hme, Avisoft), which was plugged into a computer equipped with Avisoft Recorder USG recording software (sampling frequency: 250 kHz, 16-bit format).

#### Miniature microphone

A ∼5 cm polyethylene tube (∼0.76/1.2 mm ID/OD; Cat AM801600, AM systems) was connected at one side to a 16 G blunt needle, glued to a miniature ultrasonic microphone (SPU0410LR5H-QB, Knowles), while the other side was connected to the implanted cannula before every recording session (after removal of the dummy needle from the cannula). The miniature microphone was electrically connected to the AUXin1 input of the RHD2132 amplifier board (Intan Technologies), which sampled the signals at 20 kHz. The Intan head-stage was held on the animal’s head by the Mill-Max connector glued to an Omnetics connector (NPD-36-WD-18.0-C-GS). The Intan head-stage and the miniature microphone were shielded by and grounded to an aluminum cylinder (made of a metal cigar case, 18 mm in diameter and 60 mm in length), connected to a flexible metal pipe (made of a stainless-steel shower hose, 9.5 mm in diameter and 23 cm in length; see Fig. 1A). A second shielded single-wire cable that passed through the hose electrically connected the miniature microphone to a data acquisition device (USB-1608GX, Measurement Computing), which sampled the signal at 250 kHz with a dynamic range set to ±2 V. All cables were transferred from the shower hose to their respective recording systems via a commutator (SRC-22-12V, Shenzhen Gaochong Electronics).

### Pressure recordings

To record pressure changes in the subject’s nasal cavity, we replaced the miniature microphone with a miniature pressure sensor (MB-LPS1-01-100U5N, Microbridge), connected to the Intan head-stage in an identical manner as the miniature microphone (see above). Shielding and grounding were as for the miniature microphone and relied on an aluminum cylinder (made of a metal cigar case, 19.5 mm in diameter and 82 mm in length), connected to a flexible metal pipe (made of a stainless steel shower hose, 9.5 mm in diameter and 13 cm in length). See Fig. S5.

### Recording synchronization

Recorded signals were synchronized with each other and with the video recording by a start signal generated by a custom-made triggering device (5 V TTL pulse). Both the arena microphone and video camera started recording upon this trigger signal. The same signal was recorded in the DIGin1 input of the Intan RHD2000 interface and used for synchronizing the recordings of the sniffing signals relayed to the Intan head-stage AUXin1 input from either the miniature microphone or the pressure sensor. The camera sent a TTL signal at the beginning of each frame to the Intan interface DIGin2 input, and these were used as time stamps for off-line analysis. The miniature microphone recording via the USB-1608GX data acquisition device recorded the initial trigger TTL signal as a strong pulse, enabling off-line synchronization of subsequent signals.

## QUANTIFICATION AND STATISTICAL ANALYSIS

### Tracking software

#### TrackRodent

All recorded video clips were analyzed using TrackRodent software (https://github.com/shainetser/TrackRodent), as previously described ^28^, using the *WhiteRats_TwoRatsFreeInteraction* algorithm for free interaction and the *WhiteRatWiredHeadDirectBased* algorithm for the SP task. The former algorithm uses the body contours of both animals to define video frames in which the two animals touch each other (Fig. 2A), while the latter algorithm uses the subject’s body contour to detect events when the subject investigates one of the chambers (Fig. 2B).

#### DeepLabCut

We used DeepLabCut (version 2.3.5) for multi-animal body part tracking ^32,33^. Two freely interacting rats were distinguished as “subject” or “stimulus”. We labelled eight body parts, namely left and right ears, nose, neck, trunk, left and right lateral body, and tail base (Fig. 3I), in 800 frames extracted from four videos. A ResNet-50-based neural network was employed with default parameters for 2*10^5^ training iterations. We evaluated the model with one shuffle (test error: 7.17 pixels; train error: 3.19 pixels). To improve the model, we used the outlier corrections step and retrained the model. Test error improved to 4.57 pixels and training error to 3.92 pixels. This network was then used to analyze videos from similar experimental settings. We used a p-cut-off of 0.9 likelihood to condition the X and Y coordinates for further analysis.

### Audio recording analysis

Audio signals from both microphones, sampled at 250 kHz, were analyzed first using HybrideMouse system ^56^. The system first identified the trigger pulse in the miniature microphone recording and synchronized it to the beginning of the arena microphone recording, which was started by the same trigger pulse. Next, the system used its pre-trained deep neural network model to automatically identify all USVs (defined as a discrete USV element separated from other single USV elements by at least 55 ms) separately in each recording. A manual observer then used the HybrideMouse interface to validate and correct the automatic identification, to mark each USV according to the emitter, and to identify LFVs in the same recordings. All results and audio parameters were summarized in an Excel spreadsheet (See Table S2).

### Sniffing recording analysis

The miniature microphone and pressure sensor signals recorded by the Intan head-stage were both down-sampled from 20 kHz to 5 kHz and low-pass filtered at 50 Hz. Next, we normalized the signal by subtracting the mean of the recorded signal and dividing it by its absolute maximal value. Exhalation peaks from both the microphone and the pressure sensor were determined using the Matlab function ‘islocalmin’, with the extrema detection option ‘MinProminence’ set to 0.05.

### Statistical analysis

All averaged data are shown as means ± SEM values. Statistical tests were performed using GraphPad Prism 8. The normal distribution of the data was tested using the Shapiro-Wilk tests. A paired *t*-test was used to compare different conditions or stimuli for the same single group. Two-way analysis of variance (ANOVA) tests were applied to the data to compare multiple groups and parameters. If the main effect or interaction was found, all ANOVA tests were followed by a *post hoc* Sidak’s multiple comparison test. Significance was set at 0.05 and was adjusted when multiple comparisons were used. Statistical tests regarding the number of vocalizations were done on log-transformed data. All results of the statistical analyses are detailed in Table S1.

## Supplemental Information

**Table S1**. Details of all statistical analyses, arranged according to the relevant figure (related to STAR METHODS).

**Table S2**. The processed datasets analyzed during the current study, arranged according to the relevant figure (related to STAR METHODS).

**Video S1.** A video example of a free interaction between a female subject and a female stimulus animal, together with a zoom-in on the animals, spectrograms from the arena microphone and miniature microphone, and the low-pass filtered sniffing signal from the miniature microphone (related to Figs. 1, 2, and 4).

**Video S2.** Another video example of a free interaction between a female subject and a female stimulus animal, spectrograms from the arena microphone and miniature microphone, and the low-pass filtered sniffing signal from the miniature microphone (related to Figs. 1, 2, and 4).

**Video S3.** A video example from a SP task conducted by a female subject and a male stimulus animal, together with a zoom-in on the animals, spectrograms from the arena microphone and miniature microphone, and the low-pass filtered sniffing signal from the miniature microphone (related to Figs. 1, 2, and 4).

**Video S4.** Another video example from a SP task conducted by a female subject and a male stimulus animal, together with a zoom-in on the animals, spectrograms from the arena microphone and miniature microphone, and the low-pass filtered sniffing signal from the miniature microphone (related to Figs. 1, 2, and 4).

