## Supplemental figures for "Simultaneous recording of ultrasonic vocalizations and sniffing from socially interacting individual rats using a miniature microphone"

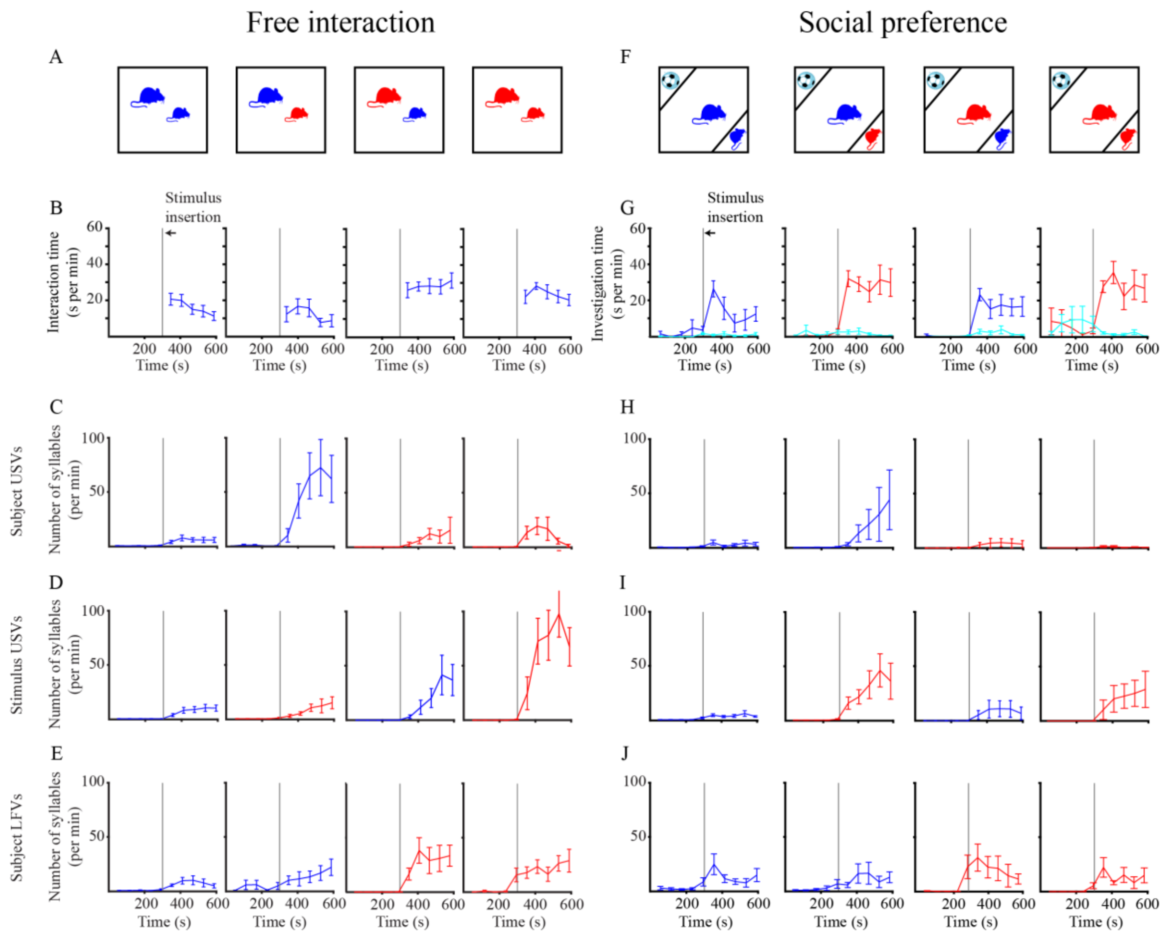


**Figure S1. Time course of social interactions and the rate of the various vocalizations in each social context (related to Fig. 3).**

- **A.** A schematic representation of the various social contexts of the free interactions sessions (blue-male, red–female, big–subject, small–stimulus animal).
- **B.** Mean (±SEM) time of physical contact (Interaction time) between the animals in each context, using 1 min bins. The grey vertical line represents the time of introduction of the stimulus animal into the arena. Note the rather constant rate of contact across the session in all cases.
- **C.** Mean (±SEM) rate of subject USVs in each context along the time course of the session, using 1 min bins. Note that subject USVs started only after stimulus introduction.
- **D.** As in **C**, for stimulus USVs. Note that stimulus USVs also started only after stimulus introduction.
- **E.** As in **C**, for subject LFVs. Note that in some cases, subject LFVs started before stimulus introduction.
- **F.** A schematic representation of the various social contexts of the SP task sessions.
- **G.** Mean (±SEM) time dedicated by subject animals during SP sessions for investigation of either the chamber of stimulus animal (blue-male, red-female) or the object's chamber (light blue) in each context, using 1 min bins. The grey vertical line represents the time of introduction of the stimulus animal into the arena. Note the clear preference for the stimulus animal over the object exhibited by the subjects in all cases.

**H-J.** As in **C-E**, for SP sessions. See also Fig. S3.

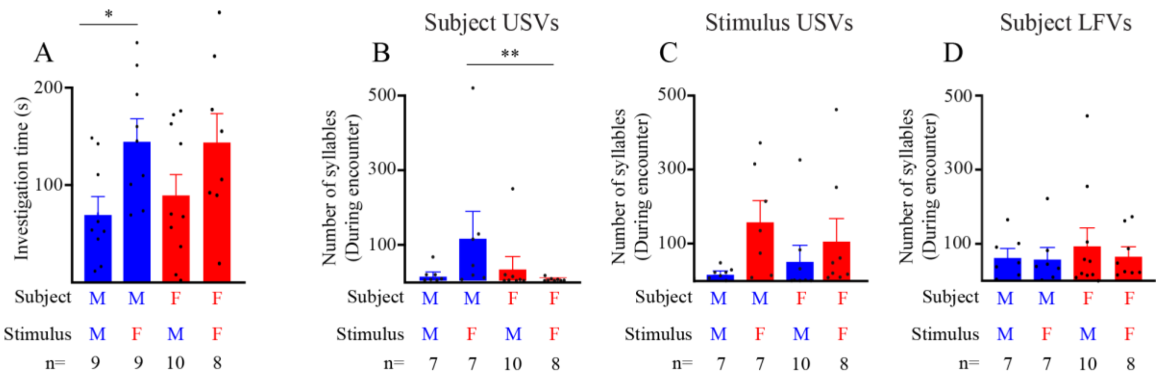


**Figure S2. The rates of the various vocalization types are sex- and social context-dependent (related to Fig. 3)**

- **A.** Mean (±SEM) social investigation time during SP sessions across the various male/female combinations (i.e, social contexts). The sex of subject and stimulus animal in each context, as well and number of animals (n), appear below. *p<0.05,
- **B.** As in **A**, for the number of subject USVs during SP sessions. **p<0.01 post hoc Sidak's multiple comparisons test following significant main effect in two-way ANOVA.
- **C.** As in **B**, for stimulus USVs.
- **D.** As in **B**, for subject LFVs.

For detailed statistical analysis results, see Table S1.

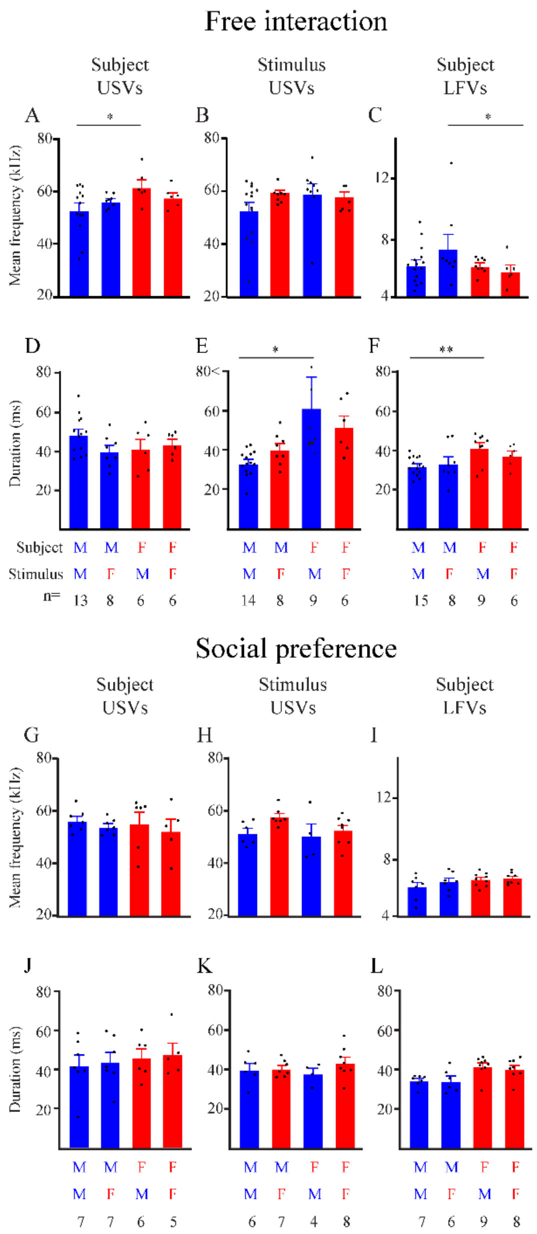


**Figure S3. Vocalization rates and spectral features are sex- and social context-dependent (related to Fig. 3)**

- **A.** Mean (±SEM) frequency of subject USVs during free social interactions in the various male/female combinations (social contexts). The sex of subject and stimulus in each context, as well and number of animals (n), appear below **D-F**.
- **B.** As in **A**, for stimulus USVs.
- **C.** As in **A**, for subject LFVs.

**D-F.** As in **A-C**, for the mean duration of the various calls.

**G-I.** As in **A-C**, for SP sessions.

**J-L.** as in **D-F**, for SP sessions.

*p<0.05, **p<0.01, ***p<0.001, according to a *post-hoc* Sidak's multiple comparison test following a significant effect in a two-way ANOVA test. For detailed statistical analysis results, see Table S1.

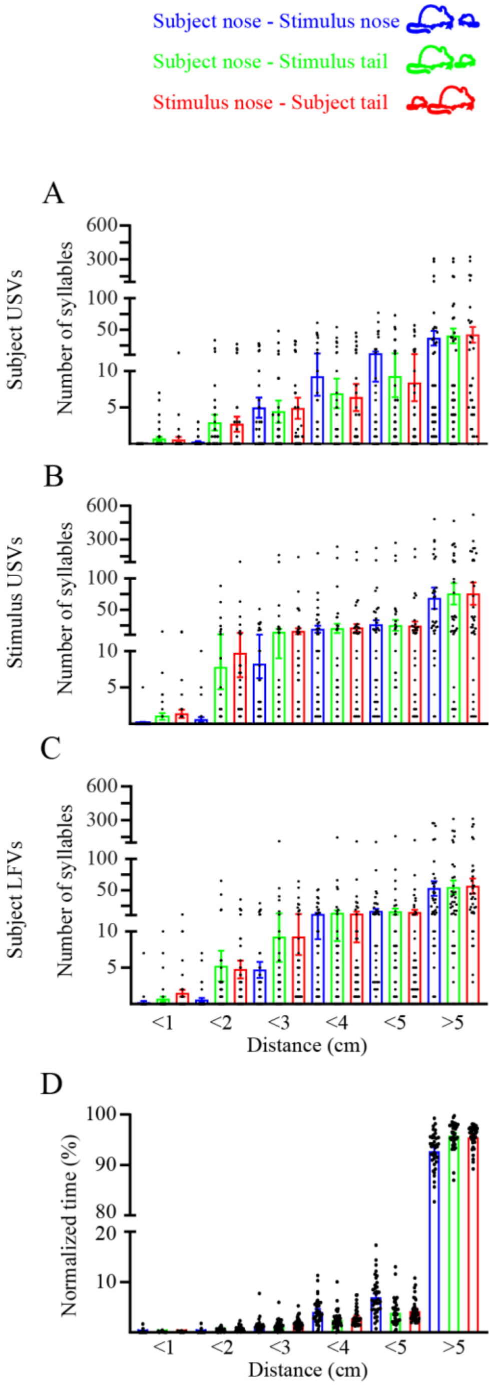


**Figure S4. Numbers of various vocalizations emitted during various behaviors and with various distances between the animals during free social interaction (related to Fig. 3)**

- **A.** Mean (±SEM) number of subject USVs emitted while the animals were in nose-to-nose (blue), nose-to-tail (green) and tail-to-nose (red) orientation (the various orientations are depicted above) at various distances from each other (see below **D**).
- **B.** As in **A**, for stimulus USVs.
- **C.** As in **A**, for subject LFVs.
- **D.** As in **A**, for the time spent in each orientation and distance, normalized to the time DeepLabCut actually detected both relevant body points.

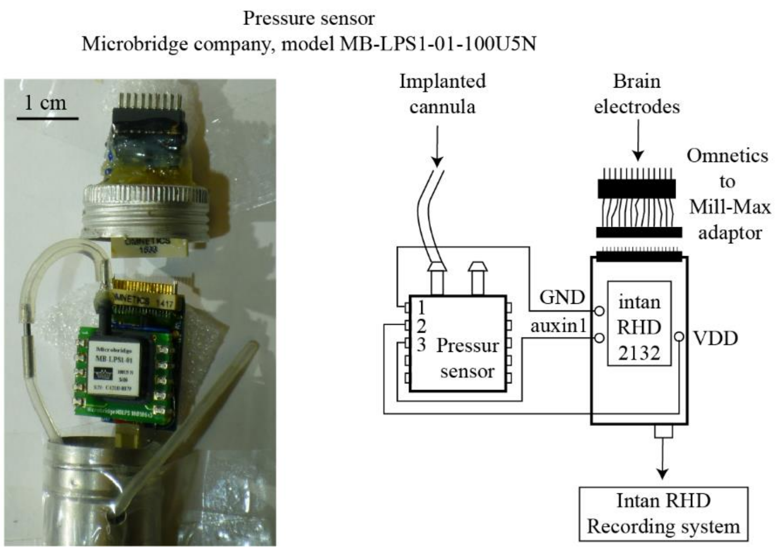


**Figure S5. The pressure sensor recording system (related to Fig. 4)**

A picture (left) and a scheme of the pressure sensor recording system, identifying the various electronic and mechanical components (see Methods).

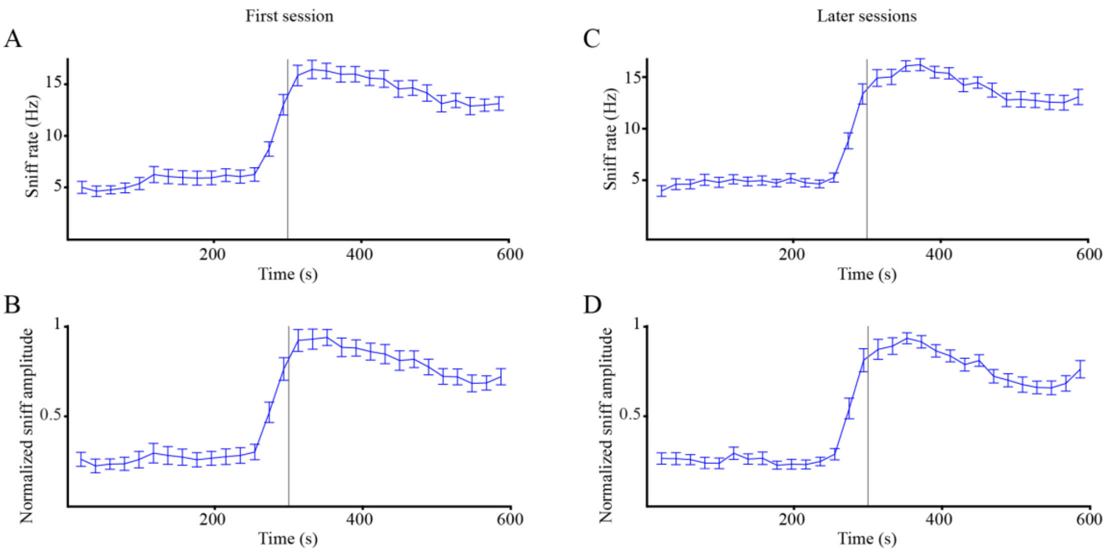


**Figure S6. Sniffing activity during the first and later sessions (related to Fig. 5)**

**A.** Mean (±SEM) sniffing rate as calculated from the miniature microphone pressure recordings in all sessions when the subject was exposed to the paradigm for the first time (first session). The vertical grey bar represents the time of stimulus introduction.

**B.** As in **A** for the normalized sniff amplitude.

**C-D**. As in **A-B**, for sessions that followed the first session (later sessions).
